# Structural-stability studies on recombinant human transferrin

**DOI:** 10.1101/742957

**Authors:** A. Kulakova, S. Indrakumar, P. Sønderby, L. Gentiluomo, W. Streicher, D. Roessner, W. Frieß, G. H.J. Peters, P. Harris

## Abstract

Transferrin is an attractive candidate for drug delivery due to its ability to cross the blood brain barrier. However, in order to be able to use it for therapeutic purposes, it is important to investigate how its stability depends on different formulation conditions. Combining high-throughput thermal and chemical denaturation studies with small angle X-ray scattering (SAXS) and molecular dynamics (MD) simulations, it was possible to connect the stability of transferrin with its conformational changes. The release of iron induces opening of transferrin, which results in a negative effect on its stability. Presence of NaCl, arginine, and histidine leads to opening of the transferrin N-lobe and has a negative impact on the overall protein stability.

**Statement of significance:** Protein-based therapeutics have become an essential part of medical treatment. They are highly specific, have high affinity and fewer off-target effects. However, stabilization of proteins is critical, time-consuming, and expensive, and it is not yet possible to predict the behavior of proteins under different conditions. The current work is focused on a molecular understanding of the stability of *human* serum transferrin; a protein which is abundant in blood serum, may pass the blood brain barrier and therefore with high potential in drug delivery. Combination of high throughput unfolding techniques and structural studies, using small angle X-ray scattering and molecular dynamic simulation, allows us to understand the behavior of transferrin on a molecular level.

## Introduction

Over the last decades, the number of approved protein-based therapeutics has increased significantly and these drugs have become essential for the treatment of various diseases, such as diabetes, hemophilia, hepatitis C, and cancer(1). This is because, compared to small molecules, protein-based therapeutics show higher specificity and therefore, generally, have less side effects. However, due to their high complexity, protein-drugs are less stable and require special conditions (formulations) that will preserve their stability during production and storage. Under inappropriate conditions, proteins have a high tendency to unfold, which may lead not only to a loss of activity, but also to aggregation(2). Unfortunately, no general rules for formulation have been reported, because it is not yet possible to predict the behavior of different proteins under different conditions. Therefore, it is important to obain a detailed molecular understanding of the rationale behind protein stability and conformational changes.

Typically the protein-drugs cannot cross the blood brain barrier (BBB), which is essential for treatment of certain diseases, such as Alzheimer’s disease and brain cancer. One of the strategies to overcome this problem is to attach protein-drugs to a protein that is able to cross the BBB. Therefore, transferrin is an attractive candidate for drug delivery(3–5), since it is one of the most abundant and stable proteins in human plasma(6), and it is able to cross the BBB through receptor-mediated endocytosis(7).

*Human* serum transferrin (TrF) is a major iron-carrying protein in the blood. TrF regulates iron levels in biological fluids, and not only supplies the cells with ferric iron, but also prevents production of radicals in the blood by removing free iron(8). TrF is a multidomain protein, which is composed of two similar lobes: the N- and the C-lobe each of them binding an iron ion hereafter refererred to iron. The lobes alter between open and closed conformation by binding and releasing iron. TrF has been crystallized in three different conformations: open(9), partially open(10), and closed(11) (see Figure 1). It has an open conformation when both lobes are free of iron. In the partially open conformation, in the presence of the transferrin receptor iron is believed to be bound to the N-lobe(12) (Fe_N_-TrF) with the C-lobe open. However, in the absence of the transferrin receptor, iron release is faster in N-lobe(12). Therefore, the crystal structure for the partially open conformation has the N-lobe open(10) with iron bound to the C-lobe(Fe_C_-TrF). Finally, TrF has a closed conformation when iron is bound to both lobes.

**Figure 1:**
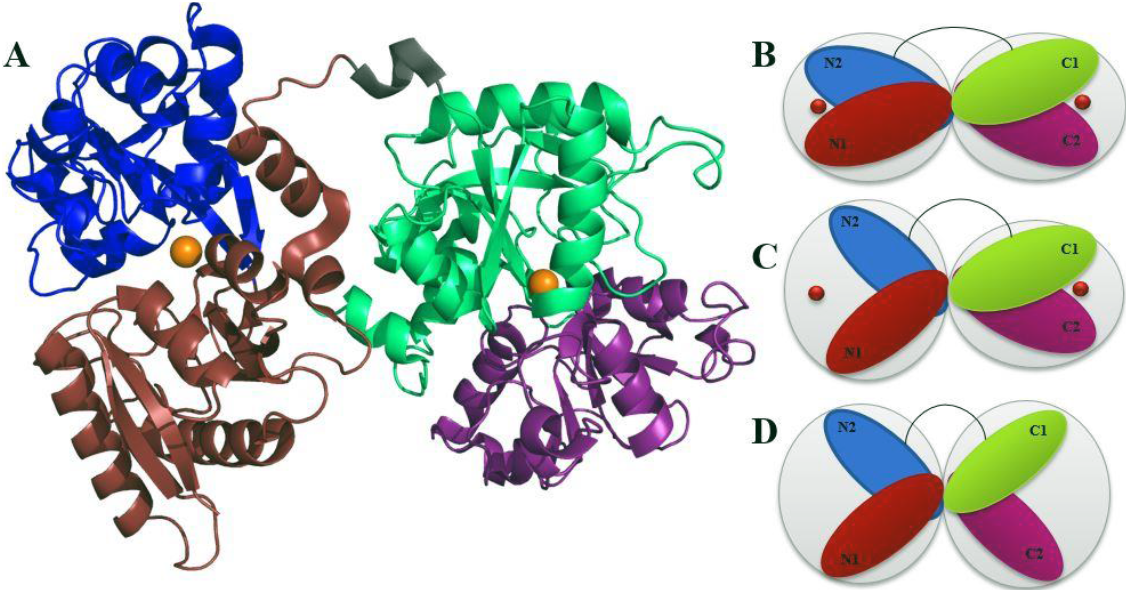
Crystal structure of the closed conformation of TrF (pdbid: 3V83)(9) and conformational representations for B: closed, C: partially open, and D: fully open conformations.

Iron release and conformational changes have been studied using a variety of techniques, including small angle scattering (SAS). Before TrF’s crystal structures became available, SAS studies indicated that TrF has spheroidal shape(13). In the presence and absence of iron, TrF showed differences in SAS curves and distance distribution functions which pointed towards conformational differences(14). Reported values for the radius of gyration were lower in the presence of iron, which suggested a more compact conformation(15–17). It has also been reported that the release of iron is pH-dependent and is induced by decreasing pH(17). Moreover, kinetic studies have shown that iron release is influenced by sodium chloride (NaCl). At neutral pH, chloride ions retards iron release, while at acidic pH, it accelerates iron release(18). In addition, it has been shown that the mechanism of iron release is a complex process that involves cooperativity between the lobes(19–21).

This study is focused on thermal and chemical denaturation of recombinant human transferrin (rTrF) in different pH and salt concentrations and with different co-solutes. These studies are combined with structural analyses performed by small-angle X-ray scattering (SAXS) and molecular dynamics (MD) simulations. The SAXS results confirmed previously reported results on the effect of pH and NaCl on the conformation: rTrF shifts towards an open conformation with decreasing pH(17) and with addition of NaCl at low pH(18), which has a negative impact on overall stability. Moreover, it was shown that arginine, which is used as common stabilizer in protein formulation, binds to rTrF destabilizing the protein as indicated by an up to 20°C decrease in the temperature of unfolding (*T*_½_). MD simulations are in agreement with the SAXS results and show that NaCl, arginine, and histidine induce opening of the N-lobe.

## Materials and Methods

### Dialysis and formulation

Recombinant human transferrin (rTrF) was provided by Albumedix Ltd. in 20 g/L solution and was dialyzed into 10 mM histidine pH 5.5, 7.0, and 10 mM tris pH 8.5 for pH and NaCl screening. Concentration of rTrF was measured on a Nanodrop™ 1000 (Thermo Fisher Scientific, Waltham, USA) using extinction coefficient calculated from the primary sequence(22) (see Table S1 in supporting information (SI)). For stability studies with different buffers and excipients, dialysis was performed at 10 mM histidine pH 5.0 and 6.5, 10 mM acetate pH 5.0 and 10 mM phosphate pH 6.5. Final solutions were obtained by diluting rTrF into the right buffer (with pH±0.1) (see Figure 2).

**Figure 2:**
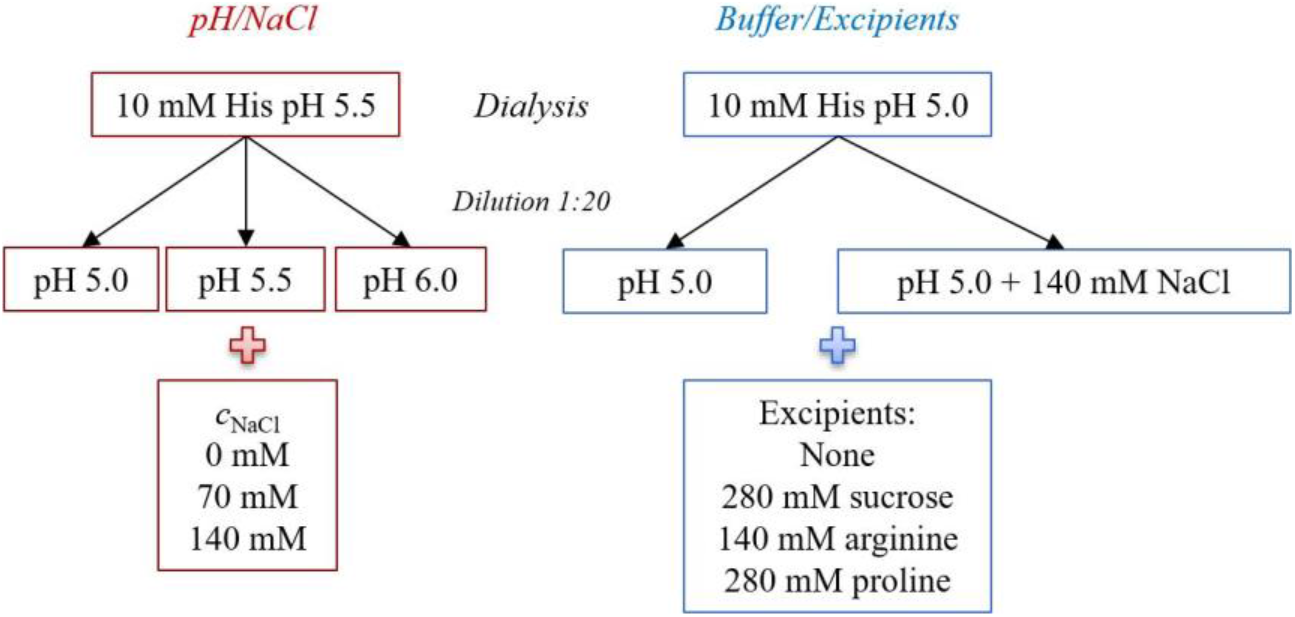
Schematic representation of dialysis and formulation process.

### Thermal stability studies

Thermal denaturation studies were performed with the Prometheus NT.48 (NanoTemper Technologies, Munich, Germany). NanoDSF Grade Standard Capillaries were manually loaded with 10 μl of protein at 1 g/L in the final conditions. All experiments were performed from 20 to 95°C with a linear thermal ramp using the heating rate of 1°C/min. Protein intrinsic fluorescence was measured and the unfolding process was monitored by looking at the shift in the fluorescence spectra (350/330 nm). All measurements were done in triplicates and the data analysis was performed using PR.Control v1.12.2 software (NanoTemper Technologies, Munich, Germany).

### Isothermal Chemical Denaturation

All chemical denaturation studies were performed on Unchained Labs HUNK system - AVIA ICD 2304 (Unchained Labs, Pleasanton, USA). The excitation wavelength was 285 nm, and emission intensities were recorded from 300 nm to 450 nm. The gain setting was set for 100, based on a previously performed gain test. From the incubation test, 162 min of additional incubation time was used. 48-point linear gradient of denaturant was automatically generated for each condition. For the first screening urea and guanidine hydrochloride (GuHCl) were used as denaturants, while for the second screening GuHCl was selected. 10 M urea and 5 M GuHCl stock solutions were prepared in each tested condition. Protein stock solutions were prepared at 1 g/L and were subsequently diluted 12.5 times to the final condition. Data collection and analysis were performed using Formulator software v3.02 (Unchained Labs, Pleasanton, USA). Protein intrinsic fluorescence was measured and the unfolding process was monitored by looking at the shift in the fluorescence spectra (356/318 nm) with increasing GuHCl concentration. In order to minimize the error, a secondary fit was performed for each pH value combining different NaCl concentrations. Free energy of unfolding (Δ*G*_unfold_), *c*_½_, and *m*-values were calculated for both transitions.

### Microscale Thermophoresis

MicroScale Thermophoresis (MST) was performed using Monolith NT.115 Label Free system through the MO.Control software (NanoTemper Technologies, Germany). All measurements were carried out in 10 mM acetate pH 5.0 at 25°C and two different ligands were chosen: arginine and proline with stock concentration of 1 M each. Each standard capillary was manually loaded with 10 μl of protein at 1 μM with different ligand concentrations, covering the concentration range from 500 to 0.78 mM. All measurement were carried out at 20% excitation power. Data analysis was done using the software MO. Affinity analysis. Initial fluorescence was used for data evaluation. For the arginine binding curve and *K*_d_ calculation eight independent experiments were performed. Proline binding affinity experiments were performed in triplicates.

### Size exclusion chromatography coupled to multi-angle light scattering (SEC-MALS)

A Vanquish Horizon™ UPLC system with a variable wavelength UV detector was operated at 280 nm (Thermo Fischer Scientific, Waltham, USA). All experiments were performed at 4 °C and temperature was controlled by autosampler. The separation was performed with a Superdex 200 increased 10/30 GL column. The aqueous mobile phase consisted of 38 mM NaH_2_PO_4_, 12 mM Na_2_HPO_4_, 150 mM NaCl and 200 ppm NaN_3_ at pH 7.4 dissolved in HPLC-grade water. The mobile phase was filtered through Durapore VVPP 0.1 m membrane filters (Millipore Corporation, Billerica, MA, USA). All the samples were centrifuged and injected in duplicates at a volume of 25 μl. Immediately after exiting the column, samples passed through the UV detector followed by static light scattering apparatus, a TREOS MALS detector (Wyatt Technology, Santa Barbara, USA), and differential refractive index detector (Optilab T-rEX, Wyatt Technology, Santa Barbara, USA). Data collection and processing were performed using the ASTRA® software V7.2 (Wyatt Technology, Santa Barbara, USA.

### Small Angle X-ray Scattering

Data collection was performed at the P12 beamline at the Petra III storage ring (DESY, Hamburg DE)(23) and at the BM29 beamline (ESRF, Grenoble FR)(24) (see Table S1 in SI). Radius of gyration (*R*_g_) and maximum dimension (*D*_max_) were derived from the experimental data with the graphical data analysis program *PRIMUSQT*(25).

The rTrF crystal structures are available in three conformations in the protein data bank(26), *i.e*, partially open (PDB ID: 3QYT(10)), closed (PDB ID: 3V83(9)), and open (PDB ID: 2HAU(11)) conformations.

Rigid body modelling of the dimer was performed using *SASREFMX*(25). In order to calculate the volume fractions of each component in the mixture, the data program *OLIGOMER*(25) was used. *FFMAKER*(25) was used to create an input file for *OLIGOMER* with a form factor for each component (open, partially open, and closed comformations retrieved from the protein data bank(26) and dimer from *SASREFMX* as input).

### Molecular Dynamics simulation

The closed rTrF crystal structure was obtained from the protein data bank(26) (PDB ID: 3V83(9)). This conformation was used as a start structure for molecular dynamics (MD) simulations. The Fe^3+^ ion and bicarbonate (CO_3_^2−^) molecules were considered during the simulations. The structure was initially prepared at pH 5.0 and pH 6.5 using the H++ server (http://biophysics.cs.vt.edu/H++)(27) which accounts for the protonation state of the titratable residues. Full details of the setup of the MD simlations has been described previously(28).The excipients acetate, phosphate, arginine, histidine, sodium chloride were included in the study. Structures were obtained from PubChem(29) and Zinc Database(30). These molecules were prepared at the desired pH using ligprep tool in Schrödinger 2016-3 suite (Schrödinger, LLC, New York, NY, USA)(31). Parameter file for the excipients and bicarbonate wereprepared using the antechamber(32) module in Amber 16 at desired pH. Charges were estimated using the AM1-BCC(33) charge method. Using the 12-6-4 LJ-type nonbonded(34, 35) model in the amber force field, parameters for Fe^3+^ were obtained. All-atom classical constant pH molecular dynamics simulations(36) in explicit solvent were carried out with the Amber 16 program(37) employing the amber force field ff99SB(38) for proteins. Titratable residues such as Asp, His, Lys, Tyr, surrounding the Fe^3+^ and the bicarbonate were titrated during the simulations. Ionic strength for each of the excipients was adjusted to 140 mM by additions of 124 solute molecules to the solvated system containing approximately 48000 water molecules. Finally, constant pH simulations were performed for 100 ns and coordinates were saved every 10 ps. Analyses were performed with CPPTRAJ(39) in Amber 16, and VMD version 1.9.3(40).

Preferential interaction coefficient (PIC) for the specific simulated system was calculated using the method described previously(28). Furthermore, an interaction score per *P*(*I*_*score*_) was calculated to estimate binding capacity of co-solute to residues on protein surface as described. Centre of mass of the co-solute was used for the determination of PIC and *P*(*I*_*score*_).

## Results

The overall conformational stability of recombinant transferrin (rTrF) was analyzed by thermal denaturation using nano differential scanning fluorimetry (nanoDSF) and isothermal chemical denaturation (ICD). Two different denaturants, urea and guanidine hydrochloride (GuHCl) were tested. Due to the high conformational stability of rTrF, urea was not strong enough to unfold it completely (see Figure 3C). Therefore, only GuHCl unfolding data were analyzed. The initial screen was performed as a function of pH (5–9) and ionic strength (0, 70, and 140 mM NaCl). NanoDSF thermal unfolding shows a single two-step transition (from folded to unfolded state), while chemical denaturation curves demonstrate the presence of an intermediate state, resulting in a three-state transition. In addition, the intermediate state is better defined at lower pH values (see Figure 3B). Only the first transition in the chemical unfolding curves was further considered, as this is where unfolding process is initiated. Due to the poorly intermediate state, the calculated Δ*G*_unfold_ shows high standard deviations and was therefore not considered for analysis.

**Figure 3:**
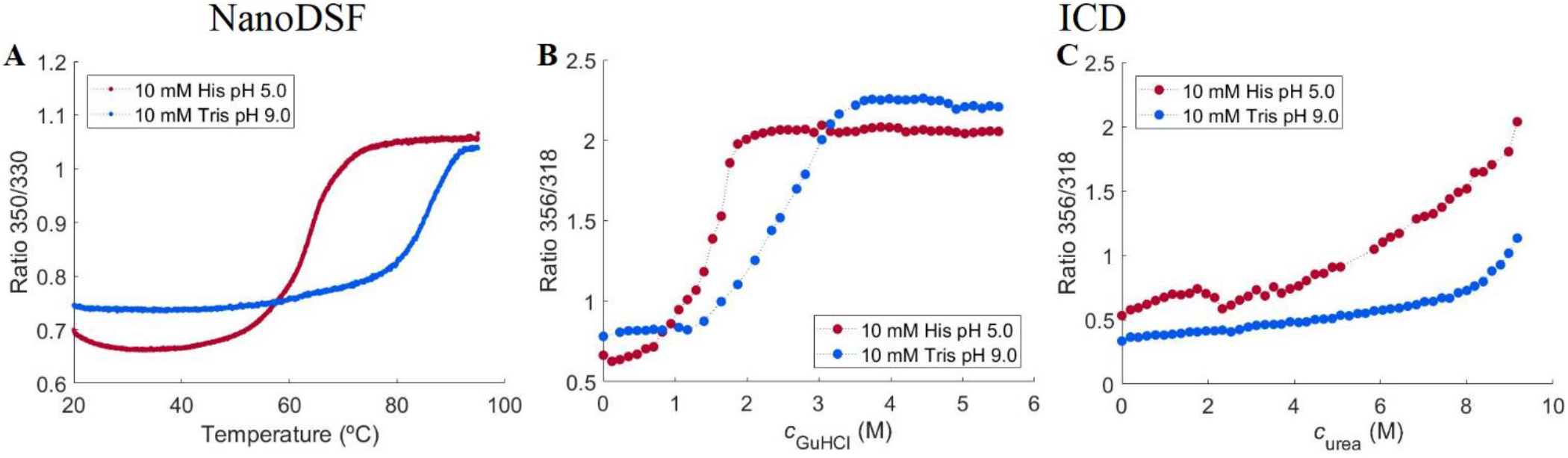
rTrF thermal and chemical unfolding curves. A: thermal unfolding curves from nanoDSF, B: chemical unfolding curves in the presence of GuHCl from ICD and C: chemical unfolding curves in the presence of urea from ICD. rTrF in 10 mM histidine (His) pH 5.0 (red), and 10 mM tris pH 9.0 (blue).

### pH dependence

*T*_½_ measured by nanoDSF is shown in Figure 4A. An increase in *T*_½_ with increasing pH is seen, meaning that the thermal stability is higher at higher pH values. Likewise, chemical denaturation shows an increase in the amount of GuHCl needed to unfold 50% of the protein (*c*_½_) with increasing pH values (see Figure 4B).

**Figure 4:**
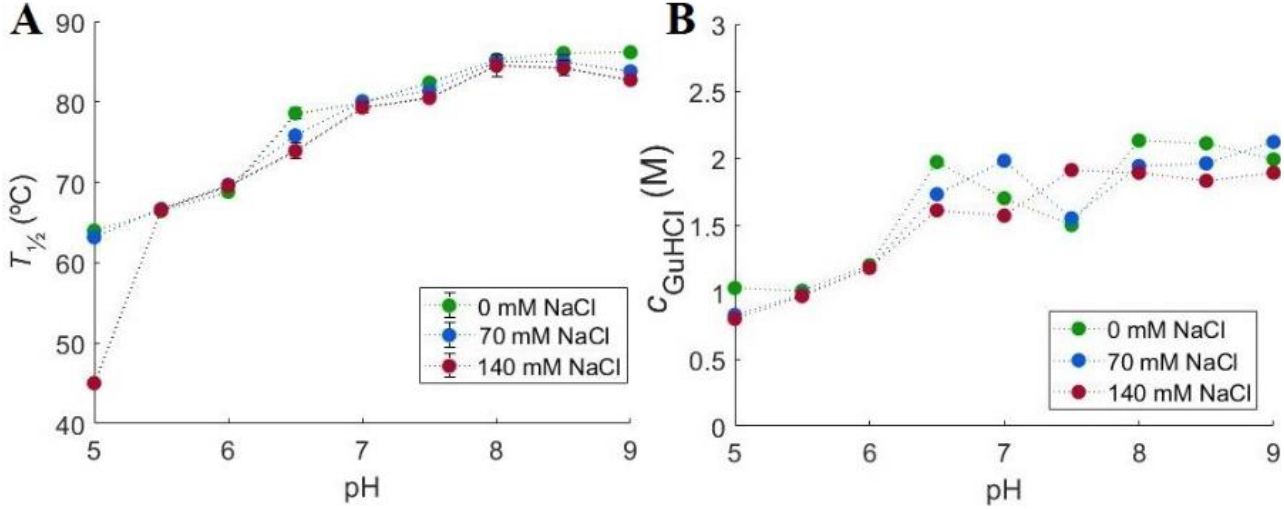
Initial stability studies performed by NanoDSF and ICD at different pH and ionic strengths. A: changes in *T*_½_ and B: changes in *c*_½_ with pH in the presence of 0 mM (green), 70 mM (blue), and 140 mM (red) NaCl.

In order to study conformational changes, SAXS concentration series data were collected at pH 4.0, 5.0, 6.5 and 8.0 with 0 mM NaCl (see Table S2 in SI). All scattering curves and SAXS data analyses are shown in the SI (see Figure S1). At pH 6.5 and 8.0, the curves coincide and the intensity at low *q*-values decreases with increasing rTrF concentration, indicating a repulsive system. Contrary to this, at pH 5.0 the intensity increases with rTrF concentration, which is characteristic for aggregation. Both aggregation and repulsion are observed at pH 4.0 (see Figure S1A in SI).

Moreover, four representative curves (shown in Figure 5) differ in shape depending on pH, which indicates conformational dissimilarity. Finally, we observe that the estimated molecular weight (*MW*) at 1 g/L for most of the conditions is higher than the expected: 75 kDa (see Table S3 in SI), which means that a substantial fraction of the protein molecules form larger species.

**Figure 5:**
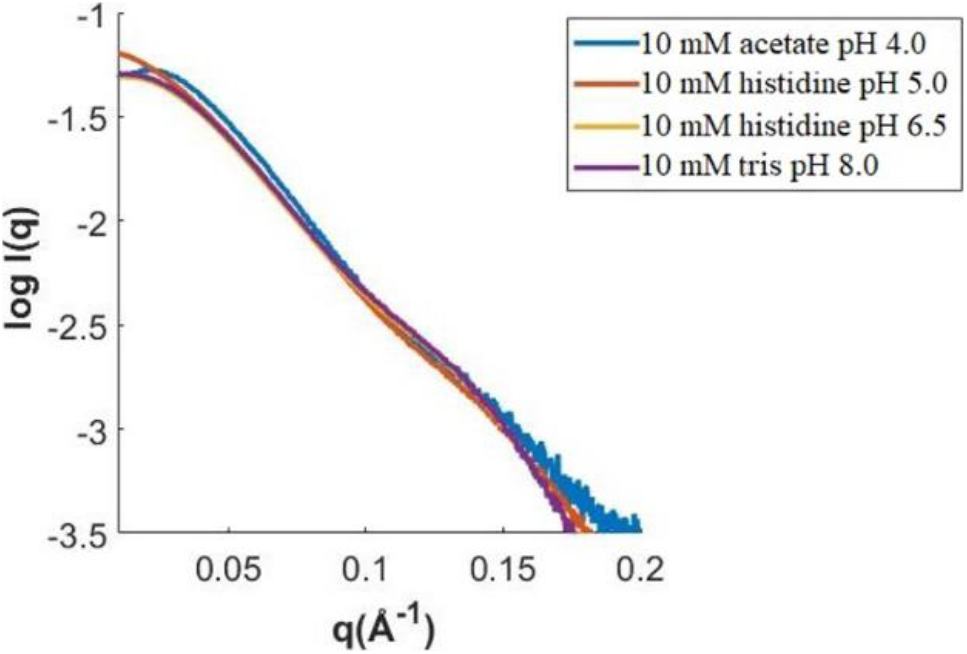
Comparison of SAXS curves from *c*_rTrF_ ~ 10 g/L collected at different pH. Blue: acetate pH 4.0; red: histidine pH 5.0; yellow: histidine pH 6.5; and purple: tris pH 8.0.

In order to characterize the size of the larger species, SEC-MALS was performed (see Table S4 and Figure S2 in SI), confirming the presence of approximately 12%, dimer and 2% trimer at all tested conditions. Additionally, static light scattering was performed as a function of protein concentration showing a concentration independent *MW* (data not shown).

It is known that rTrF exists in different conformations: open, partially open, and closed, which is related to iron binding and release(15). In order to evaluate the rTrF conformation at different pH values, *OLIGOMER*(41) analysis was performed using pdbid: 2HAU(11) for the closed conformation, pdbid: 3QYT(10) for the partially open conformation (Fe_C_-rTrF) and pdbid: 3V83(9) for the completely open conformation as input, while the dimer was modelled by *SASREFMX*(25). Due to the very small amounts of trimer, this species was not taken into consideration.

The analysis is seen in Figure 6 (see also Table S5 in SI), showing that at 10 mM histidine pH 6.5 and 10 mM tris pH 8.0 rTrF is in the closed conformation (Figure 6D and E). At 10 mM acetate pH 5.0 and 10 mM histidine pH 5.0 rTrF is present in closed and partially open conformation (Figure 6A and C).

**Figure 6:**
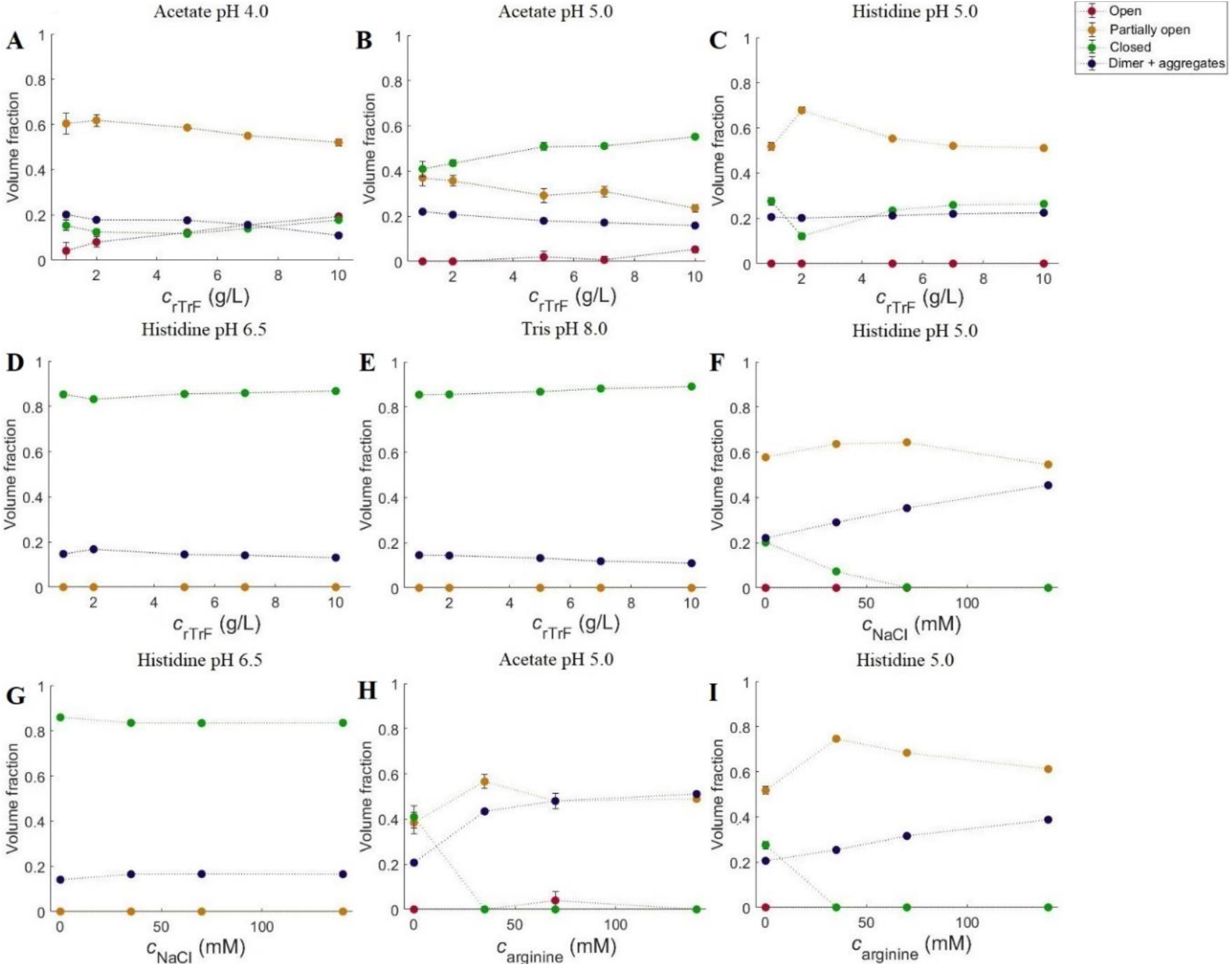
Fraction of different monomer conformations (open (in red), partially open (in orange), and closed (in green)) and dimer (in blue). A: acetate pH 4.0; B: acetate pH 5.0; C: histidine pH 5.0; D: histidine pH 6.5; E: tris pH 8.0; F: 5 g/L rTrF, histidine pH 5.0 with NaCl; G: 5 g/L rTrF, histidine 6.5 with NaCl; H: 5 g/L rTrF, acetate 5.0 with arginine; I: 5 g/L rTrF, histidine 5.0 with arginine.

By changing to 10 mM acetate buffer pH 4.0, it allowed us to detect a small amount of the open conformation. At pH 4.0, rTrF shifts from partially open to open conformation, while the fraction of the closed conformation remains constant.

### NaCl dependence

Overall thermal stability is independent of the NaCl concentration, except at pH 5.0, where 140 mM NaCl cause a decrease in *T*_½_ from 65°C to 45°C (see Figure 4A). At higher pH values the addition of NaCl does not show any effect. The ICD experiments did not show an NaCl effect on *c*_½_.

SAXS experiments were performed at 5 g/L rTrF in 10 mM histidine pH 5.0 and 6.5 with increasing NaCl concentrations. At pH 5.0, a gain in *I*(0) is seen with increasing *c*_NaCl_ due to rising *MW* (up to 100 kDa), which points to the presence of aggregates (see Table S3 in SI). The *OLIGOMER* shows an fincrease in the volume fraction of higher *MW* species (see Figure 6F). The observation of larger aggregates is in agreement with NanoDSF and ICD results showing lower conformational stability at pH 5.0 with increasing salt concentration. At pH 6.5, the repulsion decreases when NaCl is added, leading to higher *I*(0) (see Figure S1H in SI).

### Buffer and excipient dependence

For the investigations of excipient and buffer effects histidine buffer at pH 5.0 and 6.5 with 0 or 140 mM NaCl, as well as acetate pH 5.0 and phosphate pH 6.5 were selected. Furthermore, three different excipients: 280 mM sucrose, 140 mM arginine, and 280 mM proline were tested.

*T*_½_ from nanoDSF and *c*_½_ from ICD are shown in Figure 7. With respect to the buffer dependence effect, it is seen that at pH 5.0 rTrF has a 5°C higher *T*_½_ in acetate buffer compared to histidine buffer, while addition of 140 mM NaCl to the histidine buffer decreases *T*_½_ by 15°C. The ICD measurements show a somewhat different picture for histidine buffer as the addition of NaCl does not influence *c*_½_, which is already significantly lower in the histidine buffer compared to the acetate buffer.

**Figure 7:**
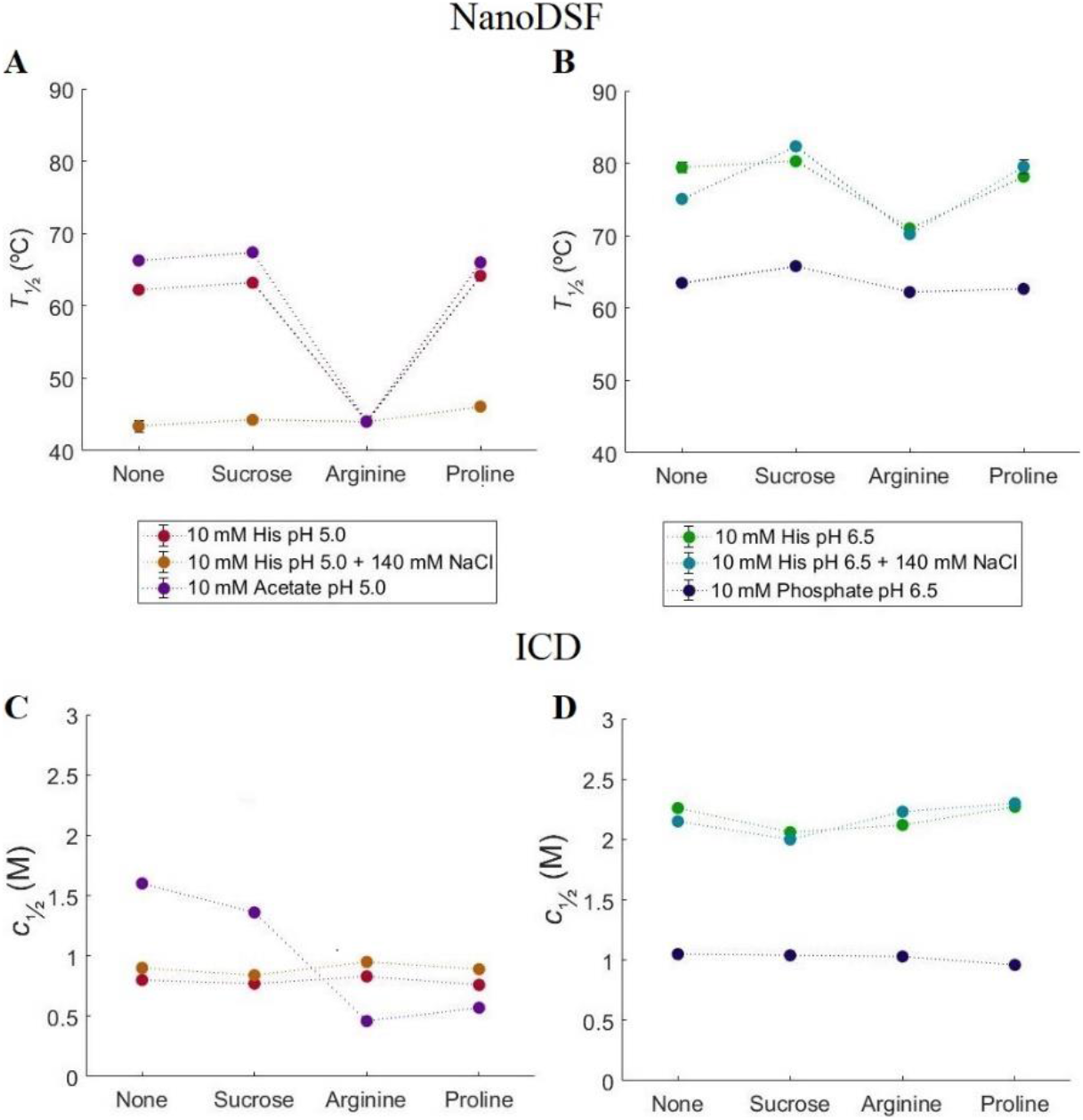
NanoDSF and ICD stability studies using different buffers and excipients. Purple: 10 mM acetate pH 5.0; red 10 mM histidine pH 5.0; orange: 10 mM histidine pH 5.0 with 140 mM NaCl; blue: 10 mM phosphate pH 6.5; green: 10 mM histidine pH 6.5; cyan: 10 mM histidine pH 6.5 with 140 mM NaCl. A: changes in *T*_½_ at histidine (0 and 140 mM NaCl) and acetate pH 5.0; B: changes in *T*_½_ at histidine (0 and 140 mM NaCl) and phosphate pH 6.5; C: changes in *c*_½_ at histidine (0 and 140 mM NaCl) and acetate pH 5.0; D: changes in *c*_½_ at histidine (0 and 140 mM NaCl) and phosphate pH 6.5.

At pH 6.5, *T*_½_ is about 15°C higher in histidine buffer than in phosphate buffer. Addition of 140 mM NaCl to the histidine buffer decreases *T*_½_ by 5°C. The ICD results are similar as *c*_½_ is reduced by 1 M in phosphate buffer, while in histidine buffer, addition of NaCl only leads to a very small decrease.

Adding sucrose or proline at pH 5.0 leads to minor effects on *T*_½._ In contrast, arginine has enormous impact on rTrF stability. Addition of arginine leads to a decrease of *T*_½_ by 20-25°C at pH 5.0 except when 140 mM NaCl is present. Both arginine and 140 mM NaCl reduce *T*_½_ by around 20°C, however, adding both of them together does not alter this already low *T*_½_ (see Figure 7A). Chemical denaturation shows a different picture, where addition of excipients in histidine buffer does not have an effect on *c*_½_. In acetate buffer *c*_½_ is 1.6 M and addition of arginine or proline leads to a decrease of *c*_½_ by approximately 1 M.

At pH 6.5 arginine decreases *T*_½_ by 5-10°C in histidine buffer, but has no effect in phosphate buffer, where *T*_½_ is already low. Excipients do not show an effect on *c*_½_ at pH 6.5 in both buffers (see Figure 7D).

The negative effect of arginine on the thermal rTrF stability can be explained by arginine binding to the protein. This was tested by performing MST using proline as a negative control, which does not have a significant effect on thermal stability. MST results show that arginine binds weakly to rTrF with *K*_d_ of 0.180 M (see Figure 8).

**Figure 8:**
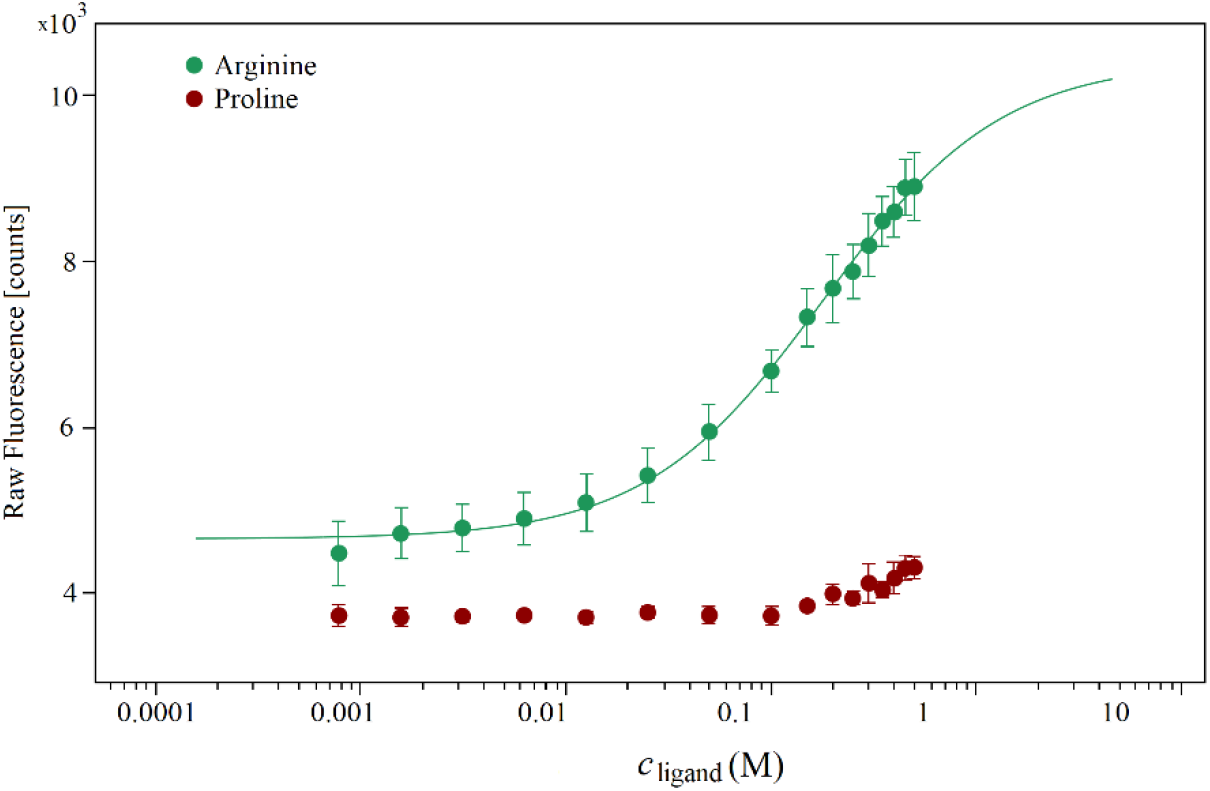
MST binding curve for arginine (in green) from *K*_d_-fit using proline (in red) as a negative control.

SAXS data were collected for rTrF in acetate and histidine buffers at pH 5.0 (see Figure S1 in SI) and analyzed using *OLIGOMER* (see Figure 6C and B, and Table S5 in SI). In both buffers, partially open and closed conformations are present, with the volume fraction of the partially open conformation decreasing with increasing rTrF concentration. However, acetate buffer shows higher volume fractions of closed conformation.

### Molecular Dynamics simulations

In order to understand the interactions between rTrF and the other components in the solution at selected pH and buffer conditions, MD simulations were performed in the presence of NaCl, histidine, arginine, acetate, and phosphate.

MD simulations were performed for 100 ns in the presence of bound Fe^3+^ and bicarbonate (CO_3_^2−^) in both lobes. All systems reached a constant root mean square deviation after 3 ns (data not shown). Patches that comprise at least three residues structurally close on the protein surface and have moderately strong interactions are colored based on *P*(*I*_*score*_) (see Figure 9). *P*(*I*_*score*_) given to a residue helps to deduct the preference of different additives on the protein surface, which in turn leads to an understanding of the different mechanism related to stabilization and iron release. At pH 5.0, both arginine and histidine are positively charged (+1), while acetate is negatively charged (−1). Even though excipients interact with the protein at several regions, only few patches interacting with the additives are relatively large and strong. Generally, arginine, histidine and NaCl bind more strongly in the C-lobe as compared to acetate, which binds stronger in the N-lobe. A particular region on the C-lobe that is a common interaction site for the different buffer components consists of residue D416 and D420 (see Figure 9A, B, and C).

**Figure 9:**
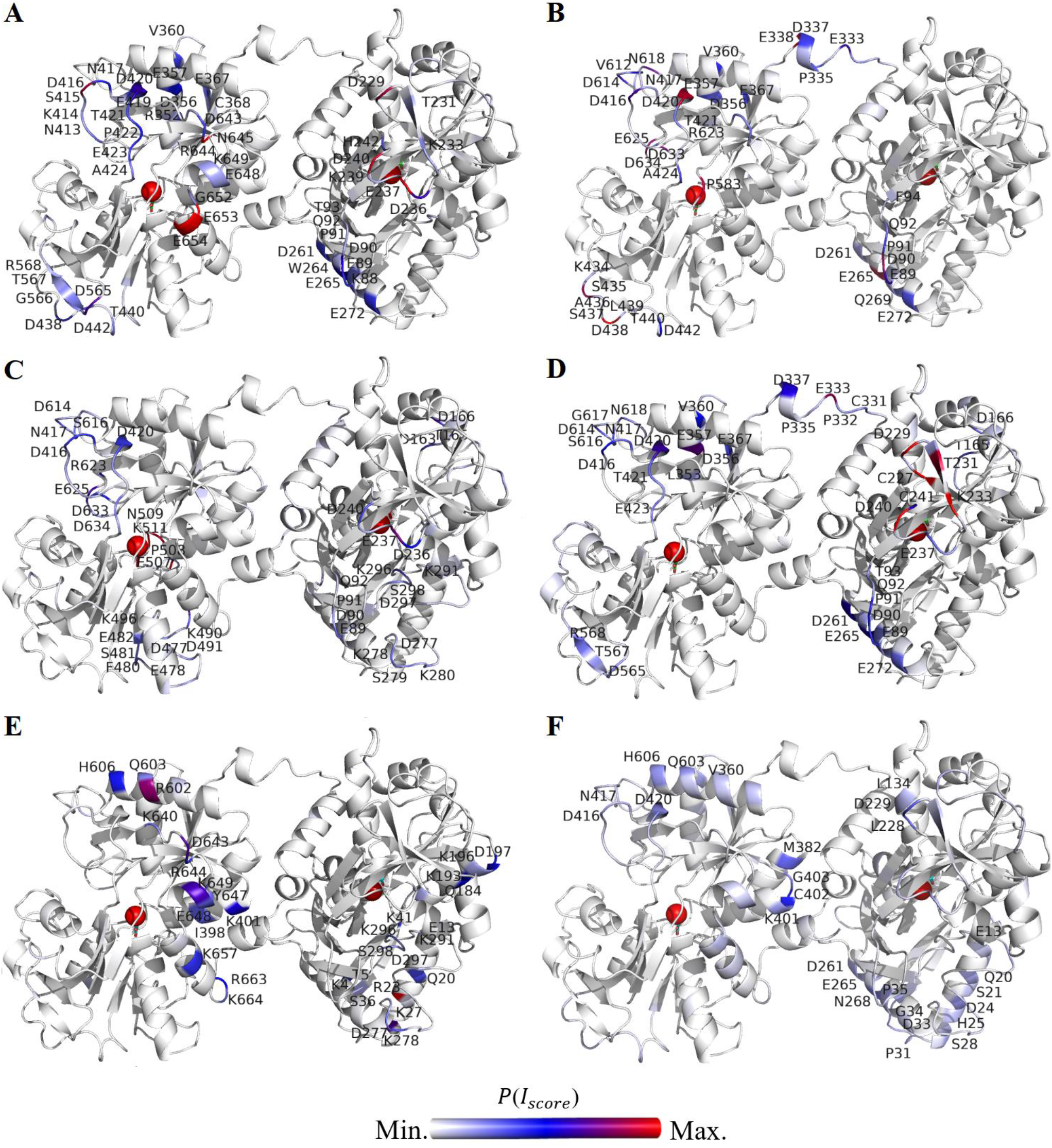
The closed structure of rTrF colored at the patches interacting strongly with buffer components based on the *P*(*I*_*score*_) calculated at pH 5.0. A: arginine, B: histidine, C: NaCl, and D: acetate; and at pH 6.5 for E: phosphate, and F: histidine. The C-lobe is shown on the left and the N-lobe on the right. Single letter code for aminoacids is used.

At pH 6.5, phosphate shows many weak interactions with single aminoacids, and relatively large, but weak interacting patches on the protein surface that are highlighted in the Figure 9E. In contrast, although interactions with histidine are spread across the protein surface, there are few prominent patches with strong interactions (see Figure 9F).

## Discussion

### pH effect

pH-dependent conformational changes in rTrF are aligned with its physiological function that is binding and transporting iron into the cells. Due to the high toxicity of iron, it is important that rTrF remains in the closed conformation in blood(8). Blood has a pH of 7.4 where according to the SAXS data only the closed conformation is present. However, rTrF should be able to supply cells with iron. Iron release occurs under acidic condition in the endosome(42, 43), where the pH is around 5(44). SAXS data analysis shows presence of the partially open conformation at pH 5.0, which supports conclusions of previous studies that prove conformational changes of rTrF being pH-dependent(17). Our SAXS studies do not show evidence of the presence of a fully open conformation at pH 5.0. At pH 4.0, a small fraction of fully open conformation was detected, accompanied by increasing aggregation (see Figure S1A in SI). This suggests that the presence of the fully open conformation may induce aggregation and therefore, it is important to have a mechanism that reduces the possibility of full opening of rTrF. This data supports that iron release is not only pH-dependent but also involves cooperativity between the N- and the C-lobe(19–21).

Thermal and chemical denaturation showed a decrease in physical stability with decreasing pH, at which the partially open conformation is present. Since this conformation has a higher solvent-accessible surface area than the closed conformation, it may unfold more easily.

### NaCl effect

With addition of NaCl, *T*_½_ at pH 5.0 decreases by 20°C and SAXS data indicate aggregation of rTrF. As already noted, at pH 5.0 both partially open and closed conformation are present. Addition of NaCl decreases the volume fraction of the closed conformation, and increases the one of the partially open conformation with the N-lobe open (see Figure 6F). Opening of the N-lobe can be explained by interactions of NaCl on the protein surface. Especially, the loop regions (residues 89-94, 277-280, and 296-298), which are close to the iron binding cleft, are prone to strongly interact with salt ions. Previous studies have shown that crosstalk between the lobes leads to iron release first occurring from the N-lobe(12, 19). At this end, strong interaction in the C-lobe loop region (D416, D420) might be inducing conformational changes that result in iron release from the N-lobe. (see Figure 9C).

This is in agreement with previous studies, where NaCl has been proven to accelerate iron release at acidic pH, due to the higher anion-binding affinity of rTrF(18) when compared to higher pH. In addition, presence of the partially open conformation, which has lower stability, contributes to the aggregation process.

At higher pH values the presence of NaCl does not induce significant changes in thermal stability. The SAXS studies confirmed that rTrF is only present in the compact conformation at pH 6.5, and that the addition of NaCl has no impact (see Figure 6G). In a previous study, chloride was shown to slow down iron release at neutral pH(18).

### Excipient effect

Amongst the tested excipients, arginine has a pronounced negative effect on the stability of rTrF, especially at pH 5.0 where *T*_½_ decreases by 20°C and *c*_½_ is reduced by 1M. According to SAXS results obtained with higher arginine concentration the fraction of partially open conformation increases at the expense of the closed conformation (see Figure 6H and I, and Table S5 in SI), leading to an aggregation. MST confirms weak arginine binding to rTrF at pH 5.0, whereas MD simulations show strong interactions in both C- and N-lobes (see Figure 9A). Adding proline in acetate has a slightly destabilizing effect seen in ICD *c*_½_, but not in *T*_½_. Furthermore, MST did not show binding of proline.

### Buffer effect

The buffer type has a clear effect on the protein stability(2). At pH 5.0, replacing histidine by acetate buffer positively affects rTrF stability, while at pH 6.5, histidine buffer is preferable over phosphate buffer.

SAXS studies show that in acetate buffer at pH 5.0 the volume fractions of the closed and partially open conformations are around 0.4 (see Figure 6B, and Table S5 in SI). In histidine buffer, the conformation of rTrF is shifted towards partially open, increasing the volume fraction to 0.5 (see Figure 6I, and Table S5 in SI). According to the MD studies, histidine has stronger binding to the C-lobe as compared to the acetate, which binds stronger in the N-lobe. However, both have a common loop region (89-94) around the iron binding site in the N-lobe, where Y96 coordinates with iron(10). Previous studies have shown that in the absence of the TrF receptor, the mechanism of iron release starts with opening of the N-lobe(12). This suggests that histidine, due to the stronger interactions with residues 89-94 in the N-lobe loop region, induces its opening and shifts equilibrium towards the partially open conformation, resulting in the lower stability.

As already mentioned, at pH 5.0, histidine, arginine, and NaCl shift the equilibrium from the closed to the partially open conformation. All of them have strong interactions with regions in the N-lobe, particularly around the iron binding site comprising of loop region (89-94), which might cause a change in electrostatic field around the region leading to conformational changes that induces the opening. Additionally, they have one common interaction patch comprising D416 and D420 and others residues around (Figure 9A, B, and C), pointing to a crucial role in rTrF’s conformational changes. These residues are present on the loop region close to C-lobe cleft, but not directly connecting the two subdomains. However, this loop is prone to high fluctuations as reflected in MD simulations (data not shown) and might be involved in the cooperativity between two lobes, since conformational changes in this region can lead to lobe-lobe interaction, and contribute to the iron release from N-lobe.

At pH 6.5 both phosphate and histidine have weak to negligible interactions in the loop region of the C-lobe and also in the loop region (89-94) of the N-lobe (see Figure 9E and F). In both buffers, rTrF is present only in the closed conformation, pointing to the involvement of these two loops (89-94, 416-420) in the iron release mechanism. Phosphate has a destabilizing effect compared to histidine, which might be due to few patches interacting strogly with phosphate on the protein surface. Contrary, histidine interacts weakly over many small patches on the protein surface. The overall charge of the phosphate at pH 6.5 is −2, while histidine is neutral, making phosphate more likely to interact strongly with the exposed hydrophilic patches as compared to histidine. Additionally, the preferential interaction coefficient values are higher for phosphate as compared to histidine implying higher preference of the protein surface for phosphate (see Figure S3 in SI).

## Conclusion

The presented work is a systematic study of the overall physical behavior of rTrF in a variety of different buffer conditions combined with structural studies using SAXS and MD simulations. The increase of *T*_½_ and *c*_½_ are both indicators of increased conformational stability. Although, some of the trends seem to be similar for these two indicators, some specific differences are seen probably because in one experiment temperature increases and in the other experiment a chemical compound is added (GuHCl). However, combining denaturation results with volume fractions of closed and partially open conformations seen in the SAXS studies (see Figure 10), it is possible to observe a decrease in volume fraction of the partially open conformation with increasing *T*_½_ and *c*_½_, and a corresponding increase in volume fraction of the closed conformation. Several conditions, such as the presence of arginine, NaCl, buffers, and pH changes can lead to opening and, consequently, to a decrease in rTrF stability. MD simulations indicate that this occurs due to the binding of the addtitives to the loop regions of the C-lobe, causing its opening for iron release.

**Figure 10:**
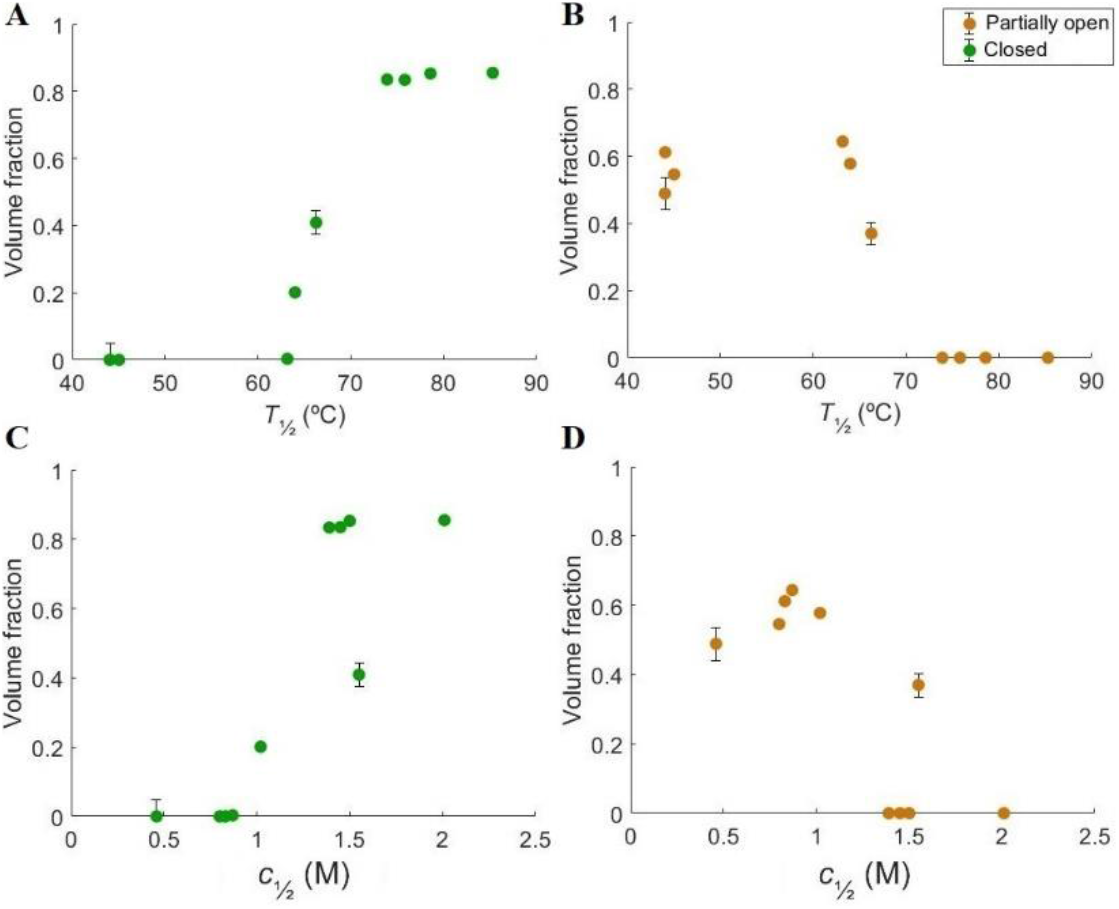
Volume fractions of different rTrF species correlated to thermal and chemical denaturation studies. A and B: volume fractions correlated to *T*_½_; C and D: volume fractions correlated to *c*_½_. Partially open conformation colored in orange and closed conformation colored in green.

## Author contributions

Alina Kulakova collected all SAXS, nanoDSF, MST and ICD data and performed formal analysis on these. She wrote original draft and edited the manuscript.

Sowmya Indrakumar performed molecular dynamic simulations and wrote that part of the manuscript.

Pernille Sønderby contributed with data interpretation for small angle x-ray scattering.

Lorenzo Gentiluomo collected SEC-malls data and performed formal analysis of these.

Günther Peters supervised MD simulation studies and participated in SAXS data collection. Pernille Harris contributed with funding acquisition, conceptualization, and supervision. Commented and edited the manuscript.

Dierk Roessner supervised LG and the light scattering experiments.

Wolfgang Friess supervised LG.

Werner Streicher supervised stability studies.

All the authors contributed with review and comments on the manuscript.

## Acknowledgements

DanScatt for funding SAXS trip.

EMBL P12 DESY and EMBL B29 ESRF for providing beam time for performing the SAXS experiments.

Albumedix Ltd for kindly providing us with recombinant transferrin.

European Union’s Horizon 2020 research and innovation program for funding (grant agreement nr 675074).

Simulations were performed at the high performance computing (HPC) services at DTU and in-house CPU/GPU cluster facilities at DTU Chemistry.

